# Risk factors for postoperative meningitis after microsurgery for vestibular schwannoma

**DOI:** 10.1101/633149

**Authors:** Bowen Huang, Yanming Ren, Chenghong Wang, Zhigang Lan, Xuhui Hui, Wenke Liu, Yuekang Zhang

**Affiliations:** Department of Neurosurgery, West China Hospital, Sichuan University, Chengdu, Sichuan, PR China

## Abstract

**OBJECTIVE:** Meningitis after microsurgery for vestibular schwannoma (VS) is a severe complication and result in high morbidity. But few studies have focused on meningitis after VS surgery alone. The purpose of this study was to identify the risk factors for meningitis after VS surgery.

**METHODS:** We undertook a retrospective analysis of all VS patients, who underwent microsurgery of VS and at least live for 7 days after surgery, between 1st June 2015 and 30st November 2018 at West China Hospital of Sichuan University. Univariate and multivariate analyses were performed to identify the risk factors for postoperative meningitis (POM).

**RESULTS:** We collected 410 patients, 27 of whom had POM. Through univariate analysis, hydrocephalus (p=0. 018), Koos grade IV (p=0.04), The operative duration (> 3 hours p=0.03) and intraoperative bleeding volume (≥ 400ml p=0. 02) were significantly correlated to POM. Multivariate analysis showed that Koos grade IV (p=0.04; OR=3.19; 95% CI 1.032-3.190), operation duration (> 3 hours p=0.03 OR= 7.927; 95% CI 1.043-60.265), and intraoperative bleeding volume (≥ 400ml p=0.02; OR=2.551; 95% CI 1.112-5.850) are the independent influencing factors of POM.

**CONCLUSIONS:** Koos grade IV, the duration of operation, and the amount of bleeding were identified as independent risk factors for POM after microsurgery of VS. POM caused a prolonged hospital stay.

## Introduction

Vestibular schwannomas (VS), also referred to as acoustic neuromas (AN), are Histopathologically benign tumors arising from Schwann cells surrounding the vestibular nerve[1], The incidence of VS is estimated to 1.9 per 100,000 per year[2], Microsurgical resection is typically the gold standard for symptomatic, relatively young patients[3], With the rapid development of minimally invasive neurosurgical technology and electrophysiological monitoring, the surgical mortality has significantly decreased[4]. However, the frequency of postoperative complications is still high. Meningitis is the main fatal complication after craniocerebral surgery. Besides, the data shows the incidence of postoperative meningitis (POM) of vestibular schwannoma surgery is about 5.5%-9.85% [4–8]. In the event of POM, the mortality can be as high as 50%[9]. However, the related research on the clinical risk factors of meningitis after acoustic neuroma surgery is limited.

Therefore, we retrospectively analyze the risk factors for meningitis following microsurgery for VS, and the frequency of this complication. The results may be able to identify the factors that significantly affect POM and provide evidence for the prevention and early clinical treatment of POM.

## Materials and methods

### Patients

This study retrospectively collected 410 patients at the Department of Neurosurgery in West China Hospital of Sichuan University, who underwent microsurgery of VS and survived at least 7 days after surgery, between 1st June 2015 and 30st November 2018. We did this study from January to February 2019. The diagnosis was based on MRI and pathology. MRI showed a mass rising from the vestibular nerve, and pathology showed schwannoma. This study was approved by the West China Hospital Ethics Committee. Written informed consent was exempted for the present study was a retrospective clinical study.

### Data collection

The basic information of these patients were collected, which include age, sex, BMI, signs and symptoms, presence/absence of diabetes mellitus, preoperative white blood cell count and hemoglobin concentration, presence/absence of hydrocephalus (assessed by magnetic resonance imaging), side and size of the tumor, history of treatment of microsurgery and/or stereotactic radiosurgery for VS, length of preoperative and postoperative hospitalization, surgery duration, bleeding amount of the operation, invasive operation, subcutaneous drainage and Lumbar drainage, Cerebrospinal fluid test data about the patients who underwent cerebrospinal fluid drainage.

### Koos grade of VS

Koos grade was utilized for the classification of VS, According to the magnetic resonance imaging of the patients and the tumor size. VS was categorized into grade I, II, III and VI.

### Surgical technique

The surgical procedures were all performed by the senior neurosurgeon (YueKang Z). Retrosigmoid approach was used for all patients. postauricular carbuncle was incised; the internal auditory canal was resected; the tumor was totally or subtotally removed by microscope; when electrophysiological monitoring was performed during the operation, the trigeminal nerve, the facial nerve, the posterior cranial nerve and the brain stem function should be perfectly protected. During the operation, prophylactic antibiotics and glucocorticoids ought to be delivered to the patients and glucocorticoids should be given continuously until the third day after the operation.

### Definition of meningitis

Meningitis must meet at least 1 of the following criteria: 1. Patient has organisms cultured from cerebrospinal fluid (CSF). 2. Patient has at least 1 of the following signs or symptoms with no other recognized cause: fever (>38°C), headache, stiff neck, meningeal signs, cranial nerve signs, or irritability and at least 1 of the following: a. increased white cells, elevated protein, and/ or decreased glucose in CSF b. organisms seen on Gram’s stain of CSF c. organisms cultured from blood d. positive antigen test of CSF, blood, or urine e. diagnostic single antibody titer (IgM) or 4-fold increase in paired sera (IgG) for pathogen[10]. These data were obtained from cerebrospinal fluid samples collected by lumbar puncture or lumbar cistern drainage.

### Statistical analysis

All the data were analyzed by using SPSS software version 25.0 (IBM Corp., Armonk, New York, USA). Continuous variables are described by median (range) and classified variables by percentages. Chi-squared test and Fisher’s exact probability test is employed to complete the univariate analysis of meningitis and non-meningitis. The factors (P <0.05) will be entered into a multivariate logistic regression analysis to determine adjusted ORs. The relationship between POM and Postoperative hospital stays will be analyzed through the Wilcoxon-Mann-Whitney test. During this analyzing process, all statistical tests were 2-sided, if P <0.05, the data will be considered to have remarkable statistics meaning.

## Results

### Baseline characteristics

In this study, 413 patients were enrolled, but three patients who died on the postoperative day were excluded. Some basic patient information was listed in table 1. The mean ages of the patients, including 172 men and 238 women, were 50 years old (range, 15-79 years). Their average BMI index was 23 Kg /m^2^(range, 16-38.6 Kg /m^2^). According to Koos, 8% of vestibular schwannomas are classified as the second level (II); 33.4% of them are classified as the third level (III); and 58.5% of them are classified as the fourth level (IV). The results show that 27 patients (6.6 %) have POM but no one died from the related meningitis.

**Table 1.** Patient characteristics and details of VS (vestibular schwannoma).

| Variables | n (%) or median (range) |
| --- | --- |
| Age | 50 (15–79) |
| Sex |  |
| Male | 172(42%) |
| Female | 238(58%) |
| BMI | 23(16–38.6) |
| Side of VS |  |
| Left | 198 (48.3%) |
| Right | 212 (51.7%) |
| Koos grade of VS |  |
| I | 0(0%) |
| II | 33 (8.0%) |
| III | 137 (33.4%) |
| IV | 240(58.5%) |

### Risk factors for POM

Through univariate analysis(Table 2), POM and hydrocephalus (p=0. 018), Koos grade IV (p=0.04), The operative duration (> 3 hours p=0.03) and intraoperative bleeding volume (≥ 400ml p=0. 02) were significantly correlated. Multivariate analysis (Table 3) showed that Koos grade IV (p0.04; OR 3.1995% CI 1.032-3.190), operation duration (> 3 hours p0.03 OR 7.92795% CI 1.043-60.265), and intraoperative bleeding volume (≥ 400ml p0.02); OR 2.551v 95% CI 1.112-5.850) are the independent influencing factors of POM.

**Table 2.**
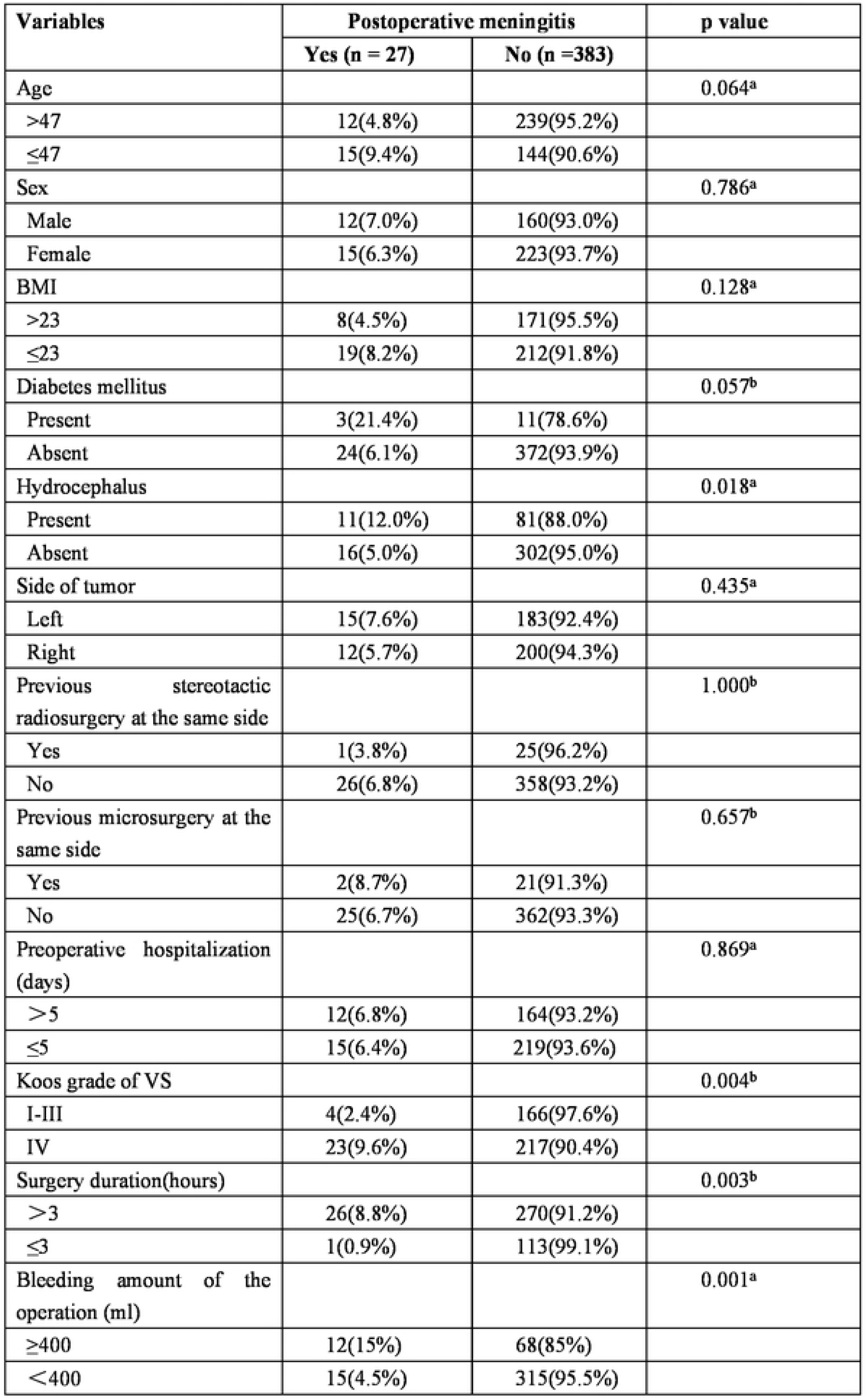
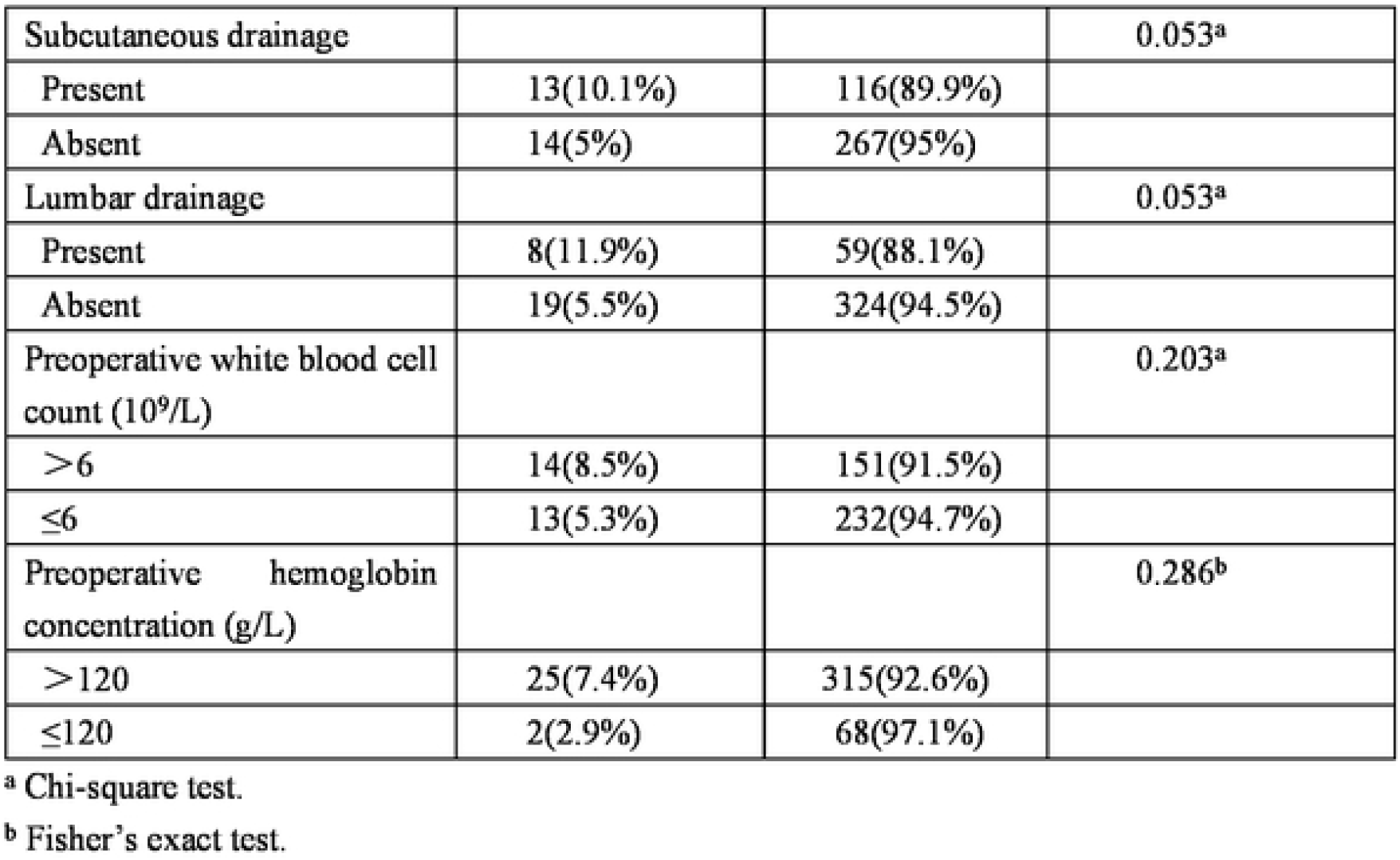
Univariate analysis of association between each factor and Postoperative meningitis.

**Table 3.** Multivariate analysis of factors associated with POM

| Variable | Odds ratio (95% CI) | p value |
| --- | --- | --- |
| Hydrocephalus (present) | 1.525(0.648-3.589) | 0.333 |
| Koos grade of VS (IV) | 3.190(1.032-9.861) | 0.044 |
| Surgery duration (>3hours) | 7.927(1.043-60.265) | 0.045 |
| Bleeding amount of the operation (≥400ml) | 2.551(1.112-5.850) | 0.027 |

### Association between POM and postoperative hospitalization

The average length of hospital stay for patients with meningitis after surgery was 16.26, with the median to be 13. The average length of hospital stay for patients without meningitis after surgery was 7.28, with the median to be 6. The result shows p < 0.001. Therefore, there was a significant difference in postoperative hospital stay between the two groups of people.

### Pathogens of POM

In this study. 27 POM patients were cultured with bacteria and fungi in CSF. The culture time was three days. However, all the culture results were negative.

## Discussion

Meningitis is divided into aseptic meningitis and bacterial meningitis. According to the previous study, the positive rate of CSF culture is about 33% [11, 12]. However, in this meningitis patient, no bacteria were cultured in the CSF culture, which may be due to the intraoperative prophylactic administration of an antibiotic, and the prophylactic administration of antibiotics in the case of postoperative infection symptoms or signs. The positive rate of CSF or blood culture decreased significantly if antibiotic is used more than 24 hours before diagnosis[13]. The culturing result turns to be negative but many cases of culture-negative (aseptic) meningitis are bacterial meningitis[14]. In this research, the incidence of meningitis after microsurgery of vestibular schwannoma is less than 6.6%, which is lower than the results reported by Huang Xiang. In their research, the incidence of meningitis after microsurgery of vestibular schwannoma is 9.85% [4]. What makes the results of the two researches inconsistent with each are as follows. First, 8% patients of ours have a tumor with the volume of less than 30 x 20 mm; Second, the data we implement in our research comes from those years from 2015 to 2018, when the surgical equipment and relevant facilities are more advanced. Third, we routinely used glucocorticoids intraoperatively and postoperatively to reduce cerebral edema and to suppress inflammatory responses. Through our data analysis, Koos grade, the duration of surgery and intraoperative blood loss are observed to be the significant factors in the development of meningitis after microsurgery of vestibular schwannoma. Besides, POM will significantly prolong the length of hospital stay.

The current research shows that the size of acoustic neuroma can significantly affect the preservation of postoperative facial nerve function[15], cerebrospinal fluid leakage[16] and postoperative pneumonia[17]. However, no literature indicates that the size of acoustic neuroma is related to POM. For the first time, our research points out that the size of the tumor significantly increased the risk of POM in the vestibular schwannoma. However, and the specific mechanism between them is still unclear. The possible mechanisms may be included as the following. First, large acoustic neuroma compress surrounding brain tissue, causing brain cells to undergo edema, damage, and inflammation; second, Grinding away the internal auditory canal produces more bone chips; third, the inflammation is caused due to the excessive traction of brain tissue during the surgery.

In addition, our study also found that the duration of surgery will significantly affect the occurrence of POM, which is consistent with previous studies. Patir et al.[18] discovered that a surgical time of more than 4 hours was significantly associated with higher postoperative infection rates. The mechanism may be that with prolonged operation time, the chance of external bacteria entering the cranium increased, and the immune function of patients was inhibited under anesthesia. Dang, Y et al.[19] claimed that Anesthesia affected the number and activity of immune cells and the secretion of cytokines. Meninges and soft tissue are retracted for surgical exposure, resulting in reduced perfusion and time-dependent reduced local immunological defense [20]. With the prolongation of the operation time, the fatigue of the operator increases, which easily pollutes the operation area.

Intraoperative hemorrhage is also an independent risk factor for meningitis in patients with vestibular schwannoma after microsurgery. Chen, C et al.[21] Chen, S et al.[22] all studied the risk factors of meningitis after microsurgery, but did not consider intraoperative hemorrhage as a risk factor. However, we found that intraoperative bleeding (> 400 ml) increased the risk of post-operative meningitis by 2.551 times. The possible reason entails further study but one possibility is that massive intraoperative hemorrhage will reduce the immune function of patients. At the same time, after massive intraoperative blood, allogeneic transfusion is often performed, but transfusion of allogeneic blood will complicate immune suppression [23]. The anti-infective ability of patients with decrease when immunity is reduced, and meningitis is prone to occur. Naidech, A. M. et al.[24] believed that the meningitis after operation was caused by subarachnoid congestion, while the blood was easily accumulated in the cerebellopontine region after vestibular schwannoma microsurgery, leading to meningitis. When patients have the above risk factors, once they have symptoms of infection, they should strongly suspect the occurrence of meningitis, and timely use effective antibiotics to control the infection.

Once meningitis occurs after surgery, many literatures reported that the hospital stay of patients with acoustic neuroma after microsurgery will be significantly prolonged[25], which is consistent with the results of this study. In addition, it will increase the medical expenses of patients, prolong the time to return to work, and increase the burden on society.

In order to prevent the complication of meningitis after the operation of acoustic neuroma, the first step is to make an early diagnosis when the acoustic neuroma is still relatively small. Larger tumors are associated with more severe symptoms and surgical complications[26]. Improving surgeon proficiency and strengthening communication and logistics among operational professionals can reduce the operation time. Golebiowski, A.[20] confirms that the duration of surgery may be shorter in older patients, so when younger patients are undergoing surgery, we should pay more attention to the control of operation time. Intraoperative hemostasis is exact, and the traction of brain tissue should be reduced. Some studies have pointed out that administration of tranexamic acid significantly reduced blood loss in patients undergoing elective craniotomy for excision of intracranial meningioma [27]. Tranexamic acid may be used to reduce intraoperative bleeding in acoustic neuroma in the future.

The disadvantage of this study is that it is a single-center study. There are admission deviations in our sample. In addition, surgical instruments and methods vary among hospitals, so our findings need to be verified in other hospitals. The average hospitalization time of our patients is only 7 days. Thus, patients suffering from meningitis after discharge are omitted from the statistics. Our study did not produce bacteria and therefore failed to analyze the distribution of meningitis bacteria. In the future, studies with a larger sample, multi-center, and more rational prospective are needed to analyze the bacterial species of meningitis in order to facilitate better treatment.

## Conclusion

In this study, the probability of meningitis after acoustic neuroma microsurgery is 6.6%. The risk factors of meningitis after microsurgery are Koos grade IV, the duration of operation (>3 hours), and the amount of bleeding (≥ 400 ml). In addition, POM has a significant association with the increase of hospitalization days after operation. Therefore, in order to prevent POM, we should shorten the operation time and reduce intraoperative bleeding.

